# Crystal structure of the human oxytocin receptor

**DOI:** 10.1101/2020.02.21.958090

**Authors:** Yann Waltenspühl, Jendrik Schöppe, Janosch Ehrenmann, Lutz Kummer, Andreas Plückthun

## Abstract

The peptide hormone oxytocin modulates socioemotional behaviour and sexual reproduction via the centrally expressed oxytocin receptor (OTR) across several species. Here, we report the crystal structure of human OTR in complex with retosiban, a non-peptide antagonist developed as an oral drug for the prevention of preterm labour. Our structure reveals insights into the detailed interactions between the G-protein coupled receptor (GPCR) and an OTR-selective antagonist. The observation of an extrahelical cholesterol molecule, binding in an unexpected location between helix IV and V, provides a structural rationale for its allosteric effect and critical influence on OTR function. Furthermore, our structure in combination with experimental data allows the identification of a conserved neurohypophyseal receptor-specific coordination site for Mg^2+^ that acts as potent positive allosteric modulator for agonist binding. Together these results further our molecular understanding of the oxytocin/vasopressin receptor family and will facilitate structure-guided development of new therapeutics.

## Introduction

Across many species, signalling mediated by oxytocin- and vasopressin-related neuropeptides provides a global, biologically conserved organising role in sexual reproduction and social behaviour^1^. In mammals, oxytocin and vasopressin orchestrate their biological role through binding and activation of closely related G protein-coupled receptors (GPCRs), called the oxytocin receptor (OTR) and the vasopressin receptors (V_1a_R, V_1b_R and V_2_R)^2,3^. The neurohypophyseal peptide hormones oxytocin and vasopressin differ only in two of their nine amino acids, but they are structurally distinct from other neuropeptides in being cyclised through a disulfide bond between two cysteines at positions 1 and 6 of the peptide chain^4,5^.

Oxytocin has numerous functions and its central neurotransmitter-like action has been shown to widely modulate cognitive effects, regulating the establishment and maintenance of complex social and bonding behaviour^6,7^, while peripheral activation of OTR is intimately linked to parturition and lactation^8–11^. Translational neuroscience research suggests that targeted activation of the OTR may have therapeutic potential in the treatment of mental health disorders including autism, Asperger’s syndrome, social anxiety disorder and schizophrenia^12–17^, while OTR antagonism can be of value in the treatment of male sexual disorders, assisted reproductive technologies and spontaneous premature labour^18–20^. However, to date, the only clinically approved OTR-targeted treatments are peptide-based ligands in obstetrics that have to be delivered through intravenous administration. To induce labour, the endogenous agonist oxytocin is administered^21^, while conversely for the prevention of preterm labour, which is still the leading cause of morbidity and mortality of children under the age of five (https://www.unicef.org/publications/index_103264.html), the antagonist atosiban is clinically approved^22^.

Despite the clear therapeutic potential of targeted intervention within this evolutionarily ancient signalling system, it has so far remained challenging to identify suitable orally available small molecule ligands that display high specificity and potency towards the OTR. Certainly, these issues can be attributed to the lack of any detailed structural information on the OTR (or the closely related vasopressin receptors) and therefore, the absence of a precise understanding of the structural determinants for ligand interaction and selectivity. Furthermore, drug discovery efforts are complicated by the OTR’s strong and incompletely understood dependence on allosteric modulators such as cholesterol and magnesium (Mg^2+^), which were shown to critically influence the interaction between the receptor and extracellular ligands^23,24^.

Here we report the first crystal structure of OTR, bound to the orally available antagonist retosiban that has been clinically developed as a tocolytic agent to efficiently block the oxytocin-mediated contraction of the smooth muscle in the uterus that occurs during the initiation of preterm labour.

In comparison to other peptidergic GPCRs, the antagonist-bound OTR structure displays an enlarged binding pocket, which is exposed to the extracellular solvent. Specific contacts with the co-crystallised antagonist retosiban are mediated through both polar and hydrophobic interacting residues that are located on opposing hemispheres of the binding cavity. Furthermore, the crystal structure allowed the identification of the binding site for the physiologically essential allosteric cholesterol molecule, which we find to be located in a pocket formed by transmembrane helices IV and V.

Finally, the identification and functional verification of two highly conserved, negatively charged residues at the extracellular tips of transmembrane helices I and II, exclusive within the neurohypophyseal oxytocin/vasopressin receptor family, allows us to postulate a structural rationale for the strong allosteric effect that Mg^2+^ exerts on agonist binding affinity. Thus, the OTR structure reported here provides the first detailed structural information on this peculiar neuropeptide-binding GPCR and provides answers to long-standing questions with regard to its allosteric modulation by both a sterol and ions.

## Results

### Structure Determination

To achieve significant purification yields of functional OTR, two consecutive rounds of directed evolution in *Saccharomyces cerevisiae* were performed on the human wild-type OTR (wtOTR)^25^. Furthermore, to allow for crystallisation in lipidic cubic phase (LCP), 34 residues (residues R232 to L265) of the third intracellular loop (ICL3) were replaced with the thermostable *Pyrococcus abysii* glycogen synthase (PGS) domain^26^, 30 residues of the flexible C-terminus (residues 360 to 389) were truncated and residues D153^4.42^ and S224^5.62^ (Ballesteros and Weinstein numbering denoted in superscript^27^) were substituted with alanine to increase thermostability. In combination, these modifications resulted in almost unaltered binding affinity for the co-crystallised antagonist retosiban (Supplementary Table 2). From crystals grown in LCP, the OTR-retosiban complex structure was determined at 3.2 Å resolution with strong and unambiguous electron density for retosiban, cholesterol and key interaction residues of the receptor (Table 1 and Supplementary Figs. 1 and 2).

**Table 1.**
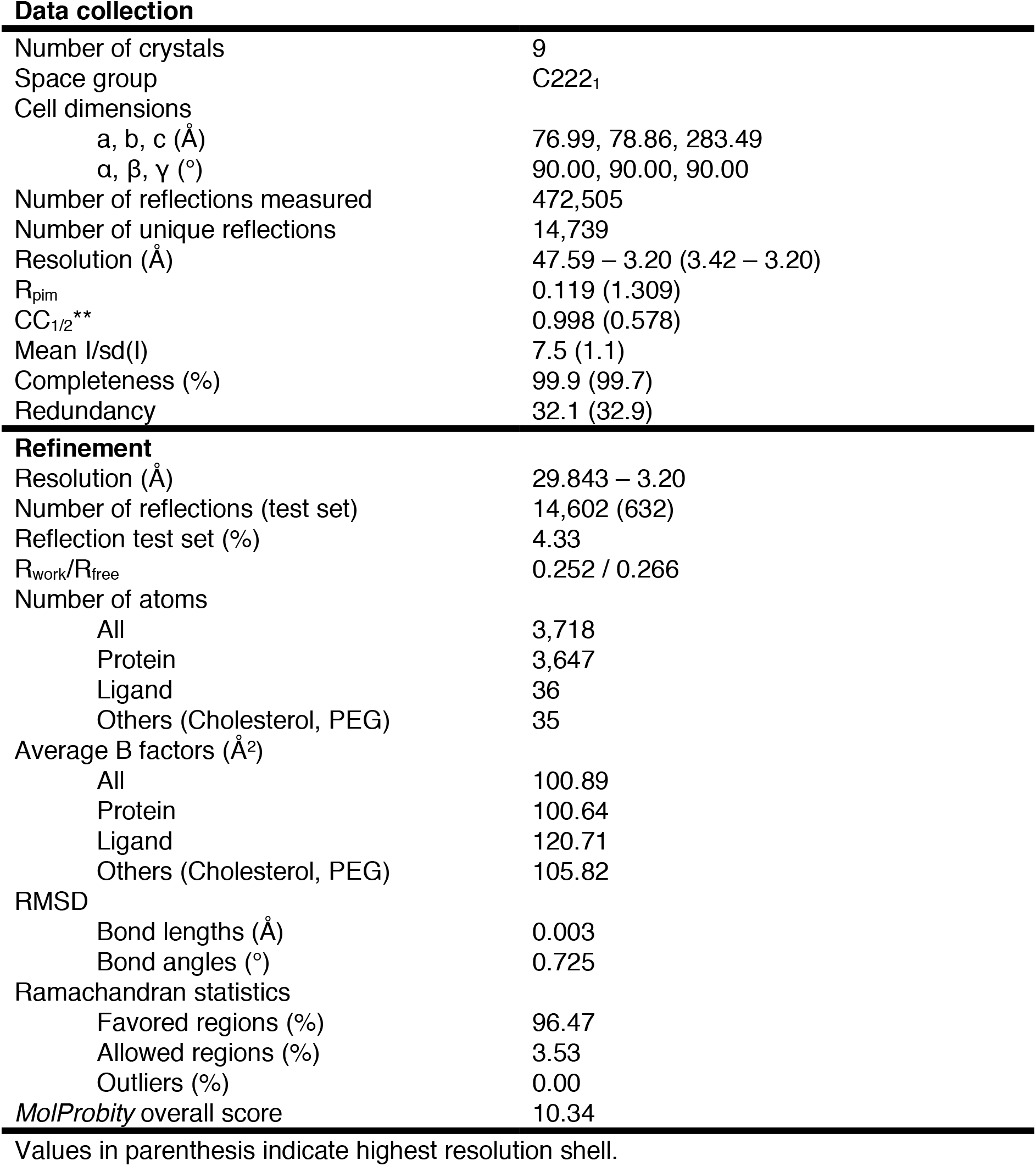
Data collection and structure refinement statistics

Overall, OTR displays the canonical GPCR topology consisting of a seven transmembrane helical bundle (helices I-VII), two intracellular loops and three extracellular loops (ICLs and ECLs, respectively) and a C-terminal amphipathic helix VIII of which only the N-terminal part is resolved (Fig. 1a,b). Similar to other class A peptide GPCRs, the ECL2 of OTR forms an extended β-hairpin structure that is anchored to the extracellular tip of helix III by the conserved disulfide bridge between C112^3.25^ and C187 of ECL2.

**Fig. 1:**
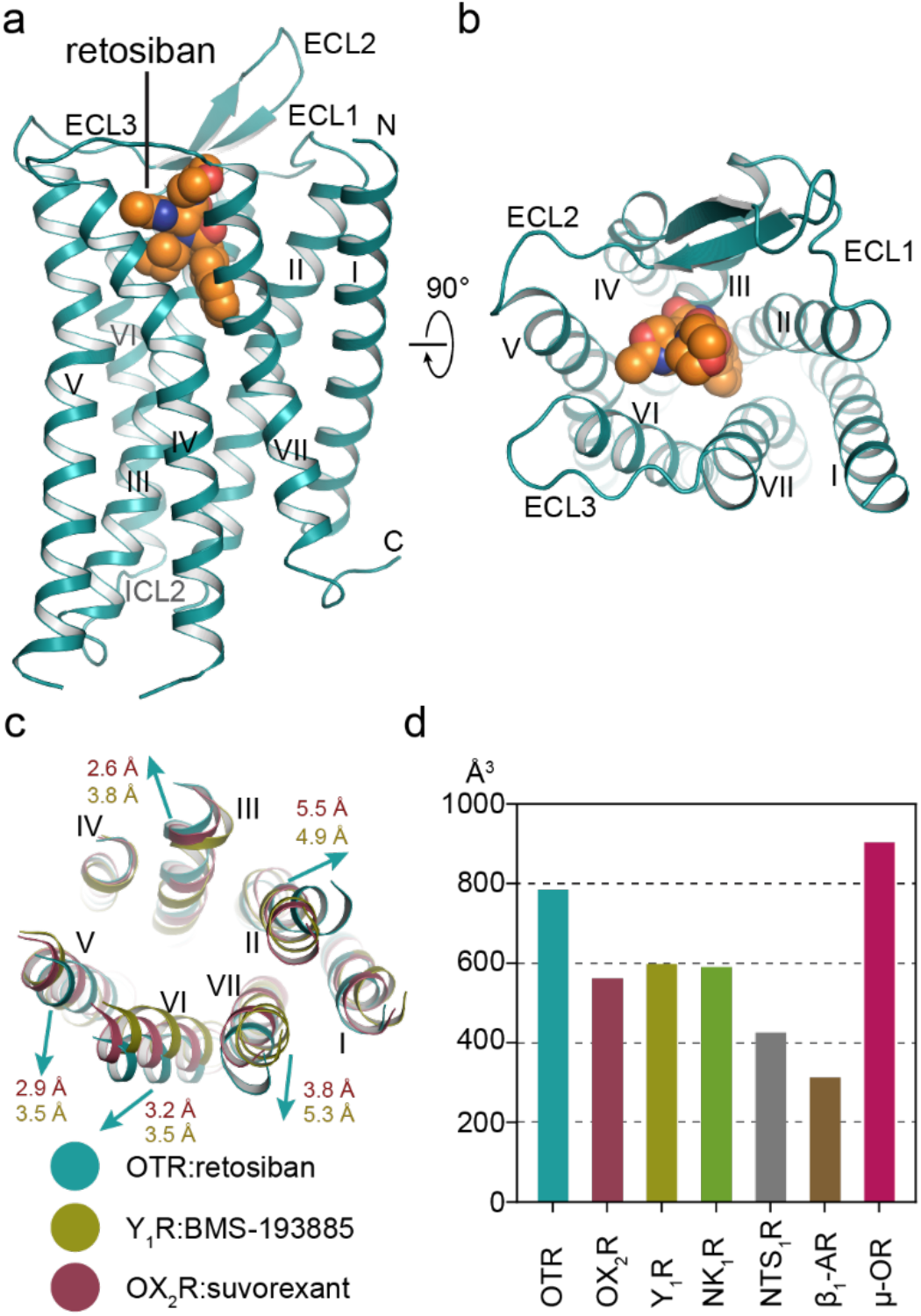
Overall structure of retosiban-bound OTR. **a**, OTR (turquoise) in complex with retosiban (orange) viewed parallel to the membrane plane. OTR is depicted as ribbon and retosiban is shown in sphere representation, with oxygen and nitrogen atoms highlighted in red and blue, respectively. **b**, OTR:retosiban complex as viewed from the extracellular space. **c**, Structural superposition of OTR with Y1R (yellow, PDB ID: 5ZBH) and OX_2_R (raspberry, PDB ID: 4S0V). Arrows indicate shifts of the OTR extracellular helix tips relative to the reference receptors. Helix tip movement in Å are indicated in the respective receptor colours (raspberry and yellow, respectively). **d**, Calculated extracellular ligand binding pocket volumes of selected neuropeptide GPCRs and one adrenergic receptor (β_1_-AR).

### Ligand-Binding Pocket of OTR

Structural superpositions with other small-molecule bound GPCRs belonging to the β-branch of class A GPCRs reveal significant conformational differences at the extracellular helical ends of OTR, even though their overall root-mean-square deviations (RMSD) for backbone atoms are as low as 1.6 Å (Supplementary Fig. 3 & Supplementary Table 1)^26,28–32^. For the OTR, we find that the extracellular tips of helices II, III, V, VI and VII are moved away from the central axis of the hepta-helical bundle, with the most dramatic outward shifts observed for the extracellular ends of helices II, VI and VII (Fig. 1c and Supplementary Fig. 3). However, these extracellular conformational differences are not transferred to the intracellular side of the helical bundle since on this part of the receptor the helical conformations coincide with those of previously determined inactive peptidergic GPCR structures, consistent with the expectation that the OTR structure in complex with the antagonist retosiban is captured in the inactive state.

To quantify the consequences of the observed extracellular helical bundle expansion on the size of the solvent-accessible ligand binding pocket, we employed an analogous strategy as recently reported for the detailed analysis of the β_1_-adrenoceptor (β_1_-AR)^33^. We calculated the accessible volumes of the ligand binding sites of the different receptors (Supplementary Table 1), and from this analysis we find the OTR binding pocket volume to be enlarged by approximately 25% compared to other non-peptide antagonist-bound neuropeptide GPCRs. Moreover, in comparison to the neurotensin receptor (NTS1R) in complex with its peptide agonist, the pocket volume of retosiban-bound OTR is increased by almost 50 %, in agreement with the reported contraction of the orthosteric pocket in the agonist-bound state of NTS_1_R^34^. Overall, we find that the OTR binding pocket volume is most similar in size and shape to the previously reported exceptionally large binding site of the μ-opioid receptor (μ-OR) (Fig. 1, Supplementary Fig. 4 and Supplementary Table 1).

The increased size of the extracellular binding pocket of OTR may indicate the necessity to accommodate a cyclic peptide, a feature that is shared within the closely related oxytocin/vasopressin receptor family. Thus this feature may provide an opportunity in future drug discovery programs targeting these receptors.

### Retosiban Binding Mode

Retosiban is a potent, non-peptide antagonist of OTR with >1400-fold selectivity over the closely related vasopressin receptors (V_1a_, V_1b_ and V_2_)^35^. Chemically, retosiban (Fig. 2a) is composed of a central 2,5-diketopiperizine (2,5-DKP) core motif with three substituents at its 3, 6 and 7-positions (butan-2-yl, indanyl and 2’-methyl-1’,3’-oxazol-4’-yl morpholine amide, respectively) that were found to provide the best balance between potency and selectivity of all tested compounds^35,36^.

**Fig. 2:**
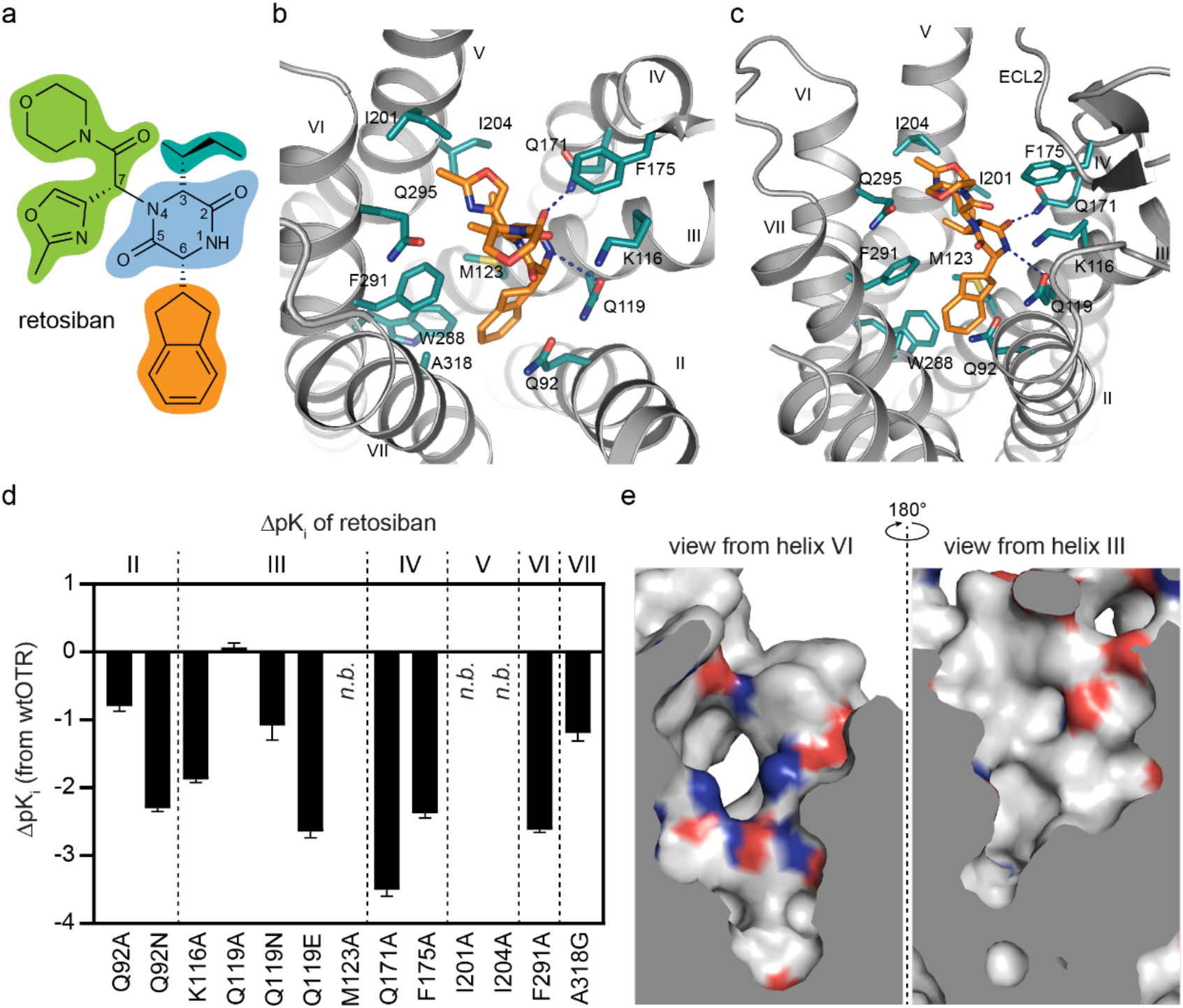
The OTR binding pocket for retosiban. **a**, Chemical structure of retosiban with structural topology highlighted by coloured circles (2,5-diketopiperazine core in blue, indanyl group in orange, sec-butyl group in cyan, oxazol-morpholine amide moiety in green). **b**, Detailed interactions of retosiban with OTR as viewed from the extracellular space from a position above helices I and II. The receptor backbone is shown in grey. Retosiban and key interaction residues within 4 Å of the ligand are shown as sticks and are coloured as in Fig. 1. Hydrogen bonds are indicated by dashed, blue lines. **c**, Interactions of retosiban with OTR viewed from the membrane-plane. d, Retosiban affinity profiles of selected mutants in comparison to wtOTR. Bars represent differences in calculated affinity (pKi) values for each mutant relative to the wild-type receptor. pKi values were derived from competition ligand binding experiments (Supplementary Table 2). Data are shown as mean values ± SEM from three independent experiments performed in quadruplicates; n.b., no binding. **e**, The two chemically distinct interaction surfaces (left: polar, charged; right: hydrophobic) of the extracellular ligand binding pocket of OTR, depicted as electrostatic surfaces, with negatively and positively charged amino acids highlighted in red and blue, respectively.

Retosiban adopts an upright, elongated conformation along the central axis of the helical bundle with the indanyl located at the bottom, the 2,5-DKP core in a central position, the *sec*butyl substituent oriented towards helix V and the oxazol-morpholine amide moiety closest to the extracellular surface (Fig. 2b,c). Overall, the binding pocket can be divided into a polar interaction surface, created by a network of residues located on helices II to IV, and a largely hydrophobic hemisphere stretching from transmembrane helices V to VII (Fig. 2e).

The central 2,5-DKP core is found in a slightly kinked conformation with its 2-keto group pointing towards helix IV and the 5-keto group oriented towards helix VII. The 2,5-DKP core specifically interacts with the receptor through a polar interaction interface, which is formed by two glutamine (Q119^3.32^ and Q171^4.60^) and one lysine (K116^3.29^) residue on the surface of helices III and IV on this side of the ligand binding pocket (Figs. 2a,b,e). Q171^4.60^ forms a strong hydrogen bond (2.5 Å distance between sidechain nitrogen and ligand oxygen) with the 2-keto oxygen, while Q119^3.32^ forms a weaker hydrogen bond with N1 of the 2,5-DKP core of retosiban (3.5 Å distance between sidechain oxygen and ligand nitrogen, Fig. 2a,b). The critical contributions of these polar interactions for the high affinity binding of retosiban to the OTR are highlighted by a severe loss in binding affinity when K116^3.29^ is mutated to alanine (75-fold), Q171^4.60^ is mutated to alanine (1600-fold), and by a significant loss in binding affinity if either the length of the sidechain is varied (Q119^3.32^N, 12-fold) or if a negative charge (Q119^3.32^E, 440-fold) is introduced at position Q119^3.32^ (Fig. 2d and Supplementary Table 2).

The hydrophobic indanyl substituent attached at position 6 of the central 2,5-DKP core, which during initial structure-activity relationship (SAR) studies was shown to increase potency ~15-fold^37^, optimally penetrates into a deep, mainly hydrophobic crevice at the bottom of the binding pocket, formed by sidechain residues of helices II, III, VI and VII. Within this pocket-extension the indanyl is laterally sandwiched between Q92^2.57^ and M123^3.36^ and forms additional hydrophobic interactions with F291^6.51^, A318^7.42^ and W288^6.48^ (Fig. 2a,b).

Additional interactions between OTR and retosiban are formed by the butan-2-yl substituent and the hydrophobic sidechains I201^5.39^, I204^5.42^ and F291^6.51^ on helices V and VI. The importance of this hydrophobic interaction surface opposite to the polar network is underscored by a 214-fold loss in binding affinity of retosiban if F291^6.51^ is mutated to alanine (Fig. 2d and Supplementary Table 2). Higher up in the binding pocket, as part of the third, most solvent exposed substituent, the oxazol moiety is oriented towards I201^5.39^ and is in close proximity to Q295^6.55^, while F175^4.64^ provides an additional hydrophobic interaction with the amide linker to the morpholine ring. In line with the extensive SAR studies performed during the development of retosiban, the morpholine ring itself has no direct interactions with the receptor^36^.

Retosiban occupies a partially overlapping binding site at the bottom of the predicted orthosteric binding pocket. Importantly, retosiban directly interacts with residues Q92^2.57^, K116^3.29^, Q119^3.32^, Q171^4.60^ and Q295^6.55^ all of which were previously described to be part of the presumed orthosteric pocket of OTR based on structural studies performed on V_1A_R^38^. Of the eleven contact residues between retosiban and OTR all but I20 1^5.39^, I204^5.42^ and A318^7.42^ are conserved in the V_1A_R receptor (V217^5.39^, M220^5.42^ and G337^7.42^ in V_1A_R, respectively). Interestingly, mutating A318^7.42^ to its homologous amino acid glycine in V_1A_R has already previously been reported to influence the selectivity profile of other non-peptide OTR antagonists (L-371,257 and L-372,622)^39^. Thus, the high selectivity of retosiban for the OTR over the V_1A_R might at least be partially rationalized by these three non-conserved interaction residues in the OTR retosiban binding pocket.

### Distinct Extrahelical Cholesterol Binding Site

Cholesterol is an abundant constituent of animal cell plasma membranes and has been shown to be intimately involved in the regulation of membrane proteins. This regulatory role of cholesterol can be achieved either indirectly by its ability to modulate the physical properties of the lipid bilayer (e.g. modulation of membrane fluidity, maintenance of lipid microdomains) or directly through specific protein-cholesterol interactions^40–42^. Modulation of protein function through cholesterol has previously been reported for a number of GPCRs including the OTR^23,43,44^, and multiple class A GPCR crystal structures have revealed direct structural evidence for specific cholesterol-GPCR interactions^45–47^, albeit at different locations in the structures.

In particular the physiological function as well as the thermostability of the OTR have been shown to be critically dependent on the presence of cholesterol. Importantly, these findings provided evidence that cholesterol or its analogues act as allosteric modulators of the OTR by shifting the receptor to a high-affinity state for both agonist and antagonist binding^23,43,48^. However, despite extensive efforts, no specific cholesterol binding site has been identified for the OTR to date.

The structure of the OTR-retosiban complex displays unambiguous extrahelical electron density for one molecule of cholesterol in a distinct extrahelical surface groove that is formed by hydrophobic residues of helices IV and V and is capped towards the extracellular space by residues of ECL2 (Fig. 3a).

**Fig. 3:**
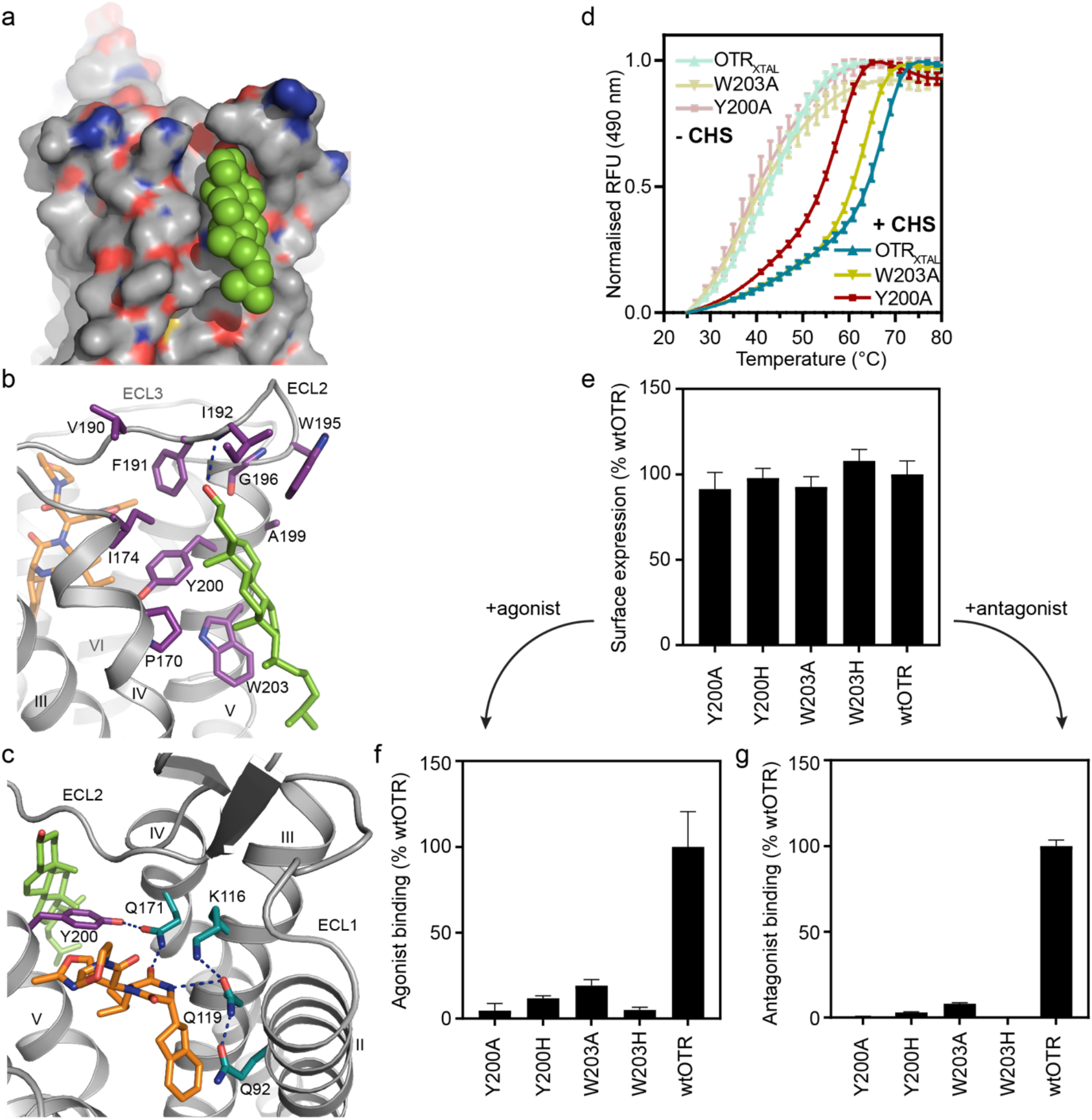
Cholesterol binding site of OTR. **a**, Surface representation of cholesterol binding site between helix IV and V of OTR, with cholesterol shown in green spheres. **b**, Detailed interactions of cholesterol with OTR, as viewed from the membrane. Residues of OTR within 4 Å of cholesterol and retosiban are depicted as sticks coloured in violet and orange, respectively. Hydrogen bonds are indicated by dashed blue lines. **c**, Cholesterol modulating the polar network of the ligand binding pocket through the positioning of Y200^5.38^, which establishes a hydrogen bond to Q171^4.60^. Through its interaction with Q171^4.60^, Y200^5.38^ coordinates the formation of a hydrogen bonding network including residues from helices II to IV. **d**, Thermostability assay (CPM) of two selected mutants in the OTR_XTAL_ background and OTR_XTAL_ performed in DDM in the presence (saturated colours) or absence (pale colours) of the cholesterol analogue cholesteryl hemisuccinate (CHS). For each experiment, a representative curve is shown. Data are shown as mean values ± SEM from 3 independent experiments. e, Receptor surface expression levels of selected mutants in wtOTR background compared to wtOTR. Data are shown as mean values ± SD from three independent experiments with 64 data points each (Supplementary Table 3). **f-g**, Maximal binding (B_max_) of agonist oxytocin (**f**) and antagonist PVA (**g**) on the OTR variants from (**e**), depicted relative to wtOTR. Data are shown as mean values ± SEM from three independent experiments in triplicates (Supplementary Table 3)

The steroid core of cholesterol specifically contacts the receptor through main hydrophobic interactions with the highly conserved (within the oxytotocin/vasopressin receptor family) residues Y200^5.38^ and W203^5.41^ (Supplementary Fig. 5) of helix V, and further with the more distal residues P170^4.59^ and I174^4.63^ on Helix IV, residues V190, F191 and W295, all of ECL2, as well as residues G196^5.43^ and A199^5.37^ on Helix V (Fig. 3b). Additionally to these hydrophobic interactions, cholesterol forms a hydrogen bond to the backbone amide of residue I192 of ECL2 with its β3-hydroxyl group.

To probe whether the observed cholesterol interaction is indeed critical for the integrity and stability of OTR, we characterised the thermal stability of purified, detergent-solubilised receptor and several mutants in the presence or absence of the cholesterol analogue cholesteryl hemisuccinate (CHS). We monitored protein unfolding using the previously described 7-diethylamino-3-(4-maleimidophenyl)-4-methylcoumarin (CPM)-based thermal shift assay^49^ (Fig. 3d). Since wtOTR could not be purified, due to very low expression yields and insufficient thermostability, the thermal stability analyses were performed with OTR_XTAL_ or mutants thereof.

In accordance with previous findings^48^, the stability of OTR_XTAL_ is significantly impaired in DDM in the absence of CHS, as evidenced by a decrease of 12°C in thermostability, when compared to the same construct measured in DDM supplemented with CHS (Fig. 3d and Supplementary Table 4). A similar effect has also been reported for several other GPCRs^50–53^. To test the individual contributions of residues within the observed cholesterol binding site, we next expressed and purified OTR_XTAL_ mutants where the two most prominent aromatic residues within the cholesterol binding site, Y200^5.38^ and W203^5.41^, were each mutated to alanine.

In the absence of CHS, OTR_XTAL_ and the two mutants Y200^5.38^A and W203^5.41^A display comparable melting temperatures (43.7°C, 44.5°C and 41.9°C, respectively), indicating a similar limited stability of all three receptor variants (Fig. 3d and Supplementary Table 4). This similarity of melting temperatures of OTR_XTAL_ mutants and OTR_XTAL_ wt shows that there is no strongly destabilising effect of the introduced mutations themselves. In the presence of CHS, in contrast, all three variants are stabilised as evidenced by higher melting temperatures, albeit to varying extents (Fig. 3d). The two tested OTR_XTAL_ mutants, where key cholesterol-interacting residues were exchanged to alanine, display a significant decrease in thermostability of 9°C (Y200^5.38^A) and 3°C (W203^5.41^A) in comparison to OTR_XTAL_, when CHS is present. This observed decrease in thermal stability of the mutants can be best rationalised by the perturbation and weakening of the binding of CHS to the cholesterol binding pocket.

To further validate the biological relevance of the observed cholesterol binding site and delineate its influence on antagonist and agonist binding in a native lipid environment, we generated point mutants of Y200^5.38^ and W203^5.41^ in the background of wtOTR and probed their involvement in cholesterol binding to the OTR in a cell-based assay. Y200^5.38^ and W203^5.41^ were individually mutated to either alanine or histidine and ligand binding was assessed using a homogeneous time-resolved fluorescence (HTRF)-based assay^54^. This experimental setup allowed us to measure simultaneously total receptor surface expression levels through the donor-only signal (E_620_) originating from the fluorescently labelled SNAP-tag and the maximum attainable response B_max_ (E_665_/E_620_) (as a measure of functional protein) at saturating concentrations of a fluorescently labelled agonist or antagonist (see Methods for details).

While all tested mutants displayed total surface expression levels similar to the wtOTR (Fig. 3e), the measured B_max_ values of the respective mutants were dramatically reduced in comparison to the wild-type receptor (Figs. 3f,g). The strong reduction in the B_max_ values suggest that, although wtOTR and the respective mutants are expressed and localised in equivalent amounts on the cellular surface, upon mutation of residues involved in the cholesterol-interaction, only a fraction of receptors remains functional with regard to both agonist and antagonist binding. Therefore, also in a native environment the mutations introduced in the cholesterol binding site appear to have a detrimental effect on the structural integrity of the extracellular binding pocket itself, with direct consequences for agonist as well as antagonist binding.

Mechanistically, the receptor modulation by the presence of cholesterol in this particular binding site might occur through the direct interaction with ECL2, which has been reported to be intimately involved in agonist binding^55^. However, cholesterol appears to also directly influence the polar network of the ligand binding pocket through the positioning of Y200^5.38^, which can then form a hydrogen bond with the carbonyl side chain of Q171^4.60^ and thereby optimally orient this residue within the extracellular binding cavity (Fig. 3c), thereby rendering the OTR in a state for optimal ligand recognition and interaction – the aforementioned high-affinity state.

### Allosteric Divalent Cation Coordination Site

GPCRs are intrinsically allosteric molecules, with many of them being negatively modulated by the monovalent cation Na^+^. High-resolution structures of several receptors have provided a structural basis for this observation: an interhelical Na^+^ binding site within the conserved water-mediated hydrogen bond network connecting helices II, III, VI and VII stabilises the inactive receptor conformation in the presence of Na^+^ and leads to a measurable decrease in agonist affinity^56^. Also for the OTR, Na^+^ has been shown to act as a negative allosteric modulator with the sodium binding pocket located in the aforementioned network in the core of the receptor^57^. In contrast to the negative allosteric effect of monovalent sodium ions, divalent cations such as Mg^2+^ (but also Ni^2+^, Zn^2+^, Mn^2+^ and Co^2+^) were shown to act as positive allosteric modulators of the OTR^24,58–60^.

In the antagonist-bound OTR structure we discovered two acidic amino acids, E42^1.35^ and D100^2.65^, located at the extracellular tips of helices I and II, which are found in unusually close proximity to each other (Fig. 4a). Both residues have previously been described to be involved in the binding of endogenous cyclic peptide agonists (E42^1.35^ in OTR for oxytocin binding; and both corresponding residues, E37^1.35^ and D95^2.65^ in V_1A_R, for AVP binding^61,62^. The short distance of only 2.8 Å between the closest oxygens of the carboxyl head groups of these two amino acids in the antagonist-bound state suggests that in this arrangement in the determined structure, the two residues share a proton. We hypothesised that, based on the charged nature and close proximity of these residues to the presumed orthosteric binding site of the OTR, this region might represent the elusive coordination site for divalent cations, known to be involved in modulating neurohypophyseal hormone binding. We note, however, that in the captured antagonist-bound state no divalent cation could be observed.

**Fig. 4:**
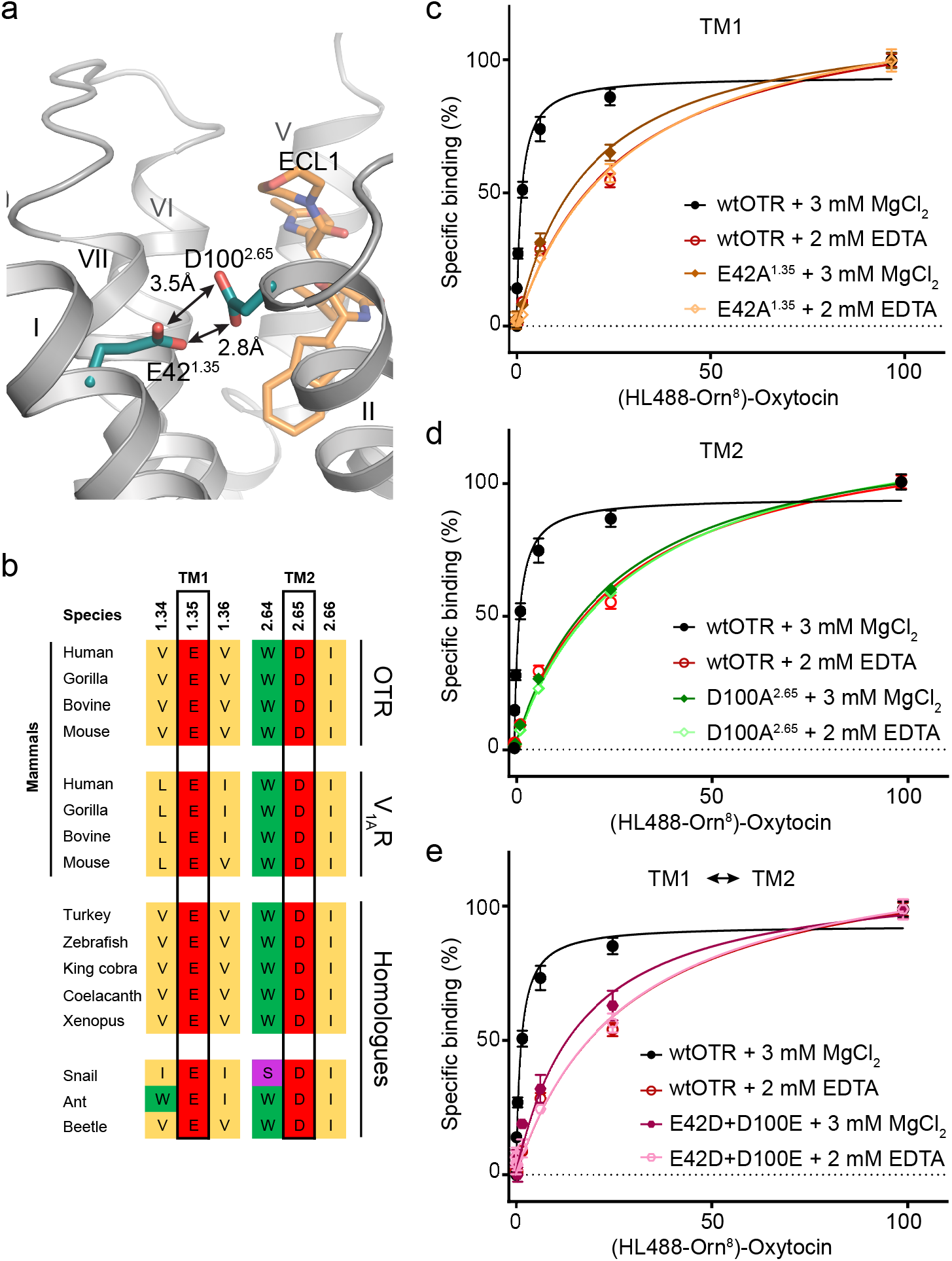
Divalent cation binding site of OTR. **a**, Location of the allosteric divalent cation binding site created by E42^1.35^ and D100^2.65^ shown in cyan sticks with head group oxygen-oxygen distances indicated by arrows. **b**, Amino acid sequence alignment of specific residues comprising the human OTR divalent cation binding site (black boxes). The divalent cation binding site of human OTR is highly conserved in homologues from different taxa including mammals, non-mammalian vertebrates and invertebrates. Furthermore, the divalent cation binding site of OTR is also conserved in mammalian homologues (human, gorilla, bovine and mouse) of the closely-related V_1A_R of the oxytocin/vasopressin system. Accession codes are provided in Supplementary Fig. 5a. **c**, Specific oxytocin binding to wtOTR and wtOTR with E42^1.35^ of the divalent cation binding site mutated to alanine (E42^1.35^A) in the presence (3 mM MgCl_2_) or absence (2 mM EDTA) of Mg^2+^. Data are shown as mean values ± SEM from three independent experiments performed in triplicates (Supplementary Table 5). **d**, As in C with the mutant D100^2.65^A. **e**, As in C with a double mutant, in which both acidic residues of the divalent cation binding site are interchanged (E42^1.35^D and D100^2.65^E).

Initial amino acid sequence alignments across both the OTR and the related V_1A_R sequences revealed that this region of transmembrane helices I and II is highly conserved in both receptors across mammalian species (Fig. 4b and Supplementary Table 4). Since the evolutionary lineage of disulfide-linked oxytocin- and vasopressin-related hormones, and thus possibly their associated GPCRs, existed for at least 700 million years^1^, we next expanded our search to receptor homologues identified in more distant taxa such as non-mammalian vertebrates and invertebrates. Indeed, we find that the glutamate in helix I and the aspartate in helix II as well as surrounding residues are strictly conserved there as well and might thus represent a mechanistically critical motif for the function of these GPCRs. Moreover, sequence comparisons with closely related class A GPCRs binding to linear peptides (Supplementary Fig. 5) did not reveal the presence of such acidic sidechains at homologous positions, thereby lending further evidence for a specific functional role of this arrangement within the neurohypophyseal hormone receptors and their related homologues.

To test if the acidic residues of helices I and II are indeed involved in the positive allosteric modulation of agonist binding through divalent cation coordination, we measured the binding affinity of fluorescently labelled oxytocin (HiLyte Fluor 488-Orn^8^ oxytocin) in the presence or absence of Mg^2+^, which is most likely the physiologically relevant cation^54^, with an HTRF-based assay. We used HEK293T cells expressing either the wtOTR, one of the single point mutants E42^1.35^A and D100^2.65^A, or the double mutant E42^1.35^D/D100^2.65^E (Fig. 4c and Supplementary Table 5). In accordance with previous studies which reported a pronounced influence of magnesium on the binding of different agonists to OTR^57,58^ we observed a 30-fold increase in affinity for oxytocin to the wild-type receptor in the presence of 3 mM Mg^2+^, compared to the same experiment performed in its absence. When divalent cations were depleted through addition of ethylenediaminetetraacetic acid (EDTA), the affinity for oxytocin to all three evaluated mutants as well as wtOTR was found to be very similar (Supplementary Table 5).

In contrast, in the presence of 3 mM Mg^2+^, the individual substitution of either E42^1.35^ or D100^2.65^ by alanine abolishes the positive modulatory effect of Mg^2+^ on oxytocin binding, thus highlighting the importance of these two specific residues in divalent cation binding. Importantly, under the same experimental conditions, the positive allosteric effect of Mg^2+^ can only be partly restored by interchanging residues, E42^1.35^D and D100^2.65^E, suggesting a significant influence of the precise spatial orientation of these two acidic residues and participation of further Mg^2+^-liganding atoms from the agonist and receptor, which is supported by the strong conservation of surrounding residues.

Taken together, these results strongly suggest that E42^1.35^ and D100^2.65^ constitute part of the physiologically important coordination site of the OTR for divalent cations and presumably also in the closely related vasopressin receptors, and that the precise spatial orientation of these residues is critical for high-affinity binding of the cyclic peptide agonists of these receptors.

## Discussion

Due to their distinct roles in social behaviour, cognition and reproduction^7^, the receptors of the oxytocin and vasopressin family represent attractive drug targets. The crystal structure of the OTR in complex with the small-molecule antagonist retosiban presented in this study provides the first structural insights into these distinct neuropeptide GPCRs; it furthermore allows us to provide answers to several long-standing questions regarding the structural basis of the essential modulatory effects by both cholesterol and divalent cations. Both of these can likely be extrapolated to the closely related vasopressin receptors due to their high sequence conservation.

The solvent-exposed binding pocket of retosiban-bound OTR is significantly larger in comparison to other antagonist-bound peptidergic GPCRs, which is possibly a direct consequence of the requirement to accommodate a cyclic nonapeptide agonist. Within the binding pocket the ligand-accessible interface, ranging from helix II to helix IV, is dominated by a polar network, while the opposing side of the cavity is largely hydrophobic. By virtue of its 2,5-diketopiperizine core and its chemically diverse substituents, the co-crystallised nonpeptide oral antagonist retosiban optimally addresses this challenging environment, lending credit to decades of SAR studies to optimise such compounds.

Additionally, the OTR structure provides new structural insights into the binding location of the modulatory cholesterol molecule. We find that the cholesterol molecule binds to an extrahelical crevice (formed between helices IV, V and ECL2) that is distal from previously reported cholesterol interaction sites on other GPCRs, consistent with non-conserved sequences in these regions^47^. Through mutagenesis experiments using orthogonal assays we established that binding of cholesterol in OTR is critically governed by the conserved aromatic residues Y200^5.38^ and W203^5.41^ on helix V. The presence of cholesterol within this binding site has a direct effect on the integrity of the binding pocket as evidenced by an almost complete loss in measurable ligand binding for antagonist as well as the agonist, if single point mutations are introduced.

This finding, obtained both from purified protein preparations *in vitro* and from cellular *in vivo* experiments, makes it conceivable to now specifically target this extrahelical binding site for the development of potent, positive or negative allosteric modulators for the OTR.

Finally, a mechanistic basis of the allosteric modulation of OTR function through interaction with Mg^2+^ is provided by the identification of an evolutionarily highly conserved Mg^2+^ binding site which is formed by two acidic residues at the extracellular tips of helices I and II, likely involving other coordinating atoms in agonist and receptor. To our knowledge, the identification and verification of this distinct, solvent-exposed coordination site in the OTR represents the first example for a positive allosteric modulatory mechanism of GPCRs by divalent cations and thus further expands our knowledge of this intrinsically allosteric receptor superfamily.

Despite their diverse physiological roles, the receptors for the cyclic peptide hormones oxytocin and vasopressin are very closely related. Thus, the studies presented here provide not only answers concerning the OTR itself, but these findings will support research advances within the vasopressin receptors, or possibly can even be directly extrapolated. We anticipate that the inaugural structure from this medically highly relevant receptor family branch opens up attractive possibilities to deploy structure-based drug-design methods to not only the OTR but eventually also to the vasopressin receptors in an effort to develop new therapeutics for the efficient treatments of a diverse and important array of diseases.

## Methods

### Generation of OTR crystallisation construct

The human OTR gene was codon-optimised for *Spodoptera frugiperda (Sf9)* expression, synthesised (Integrated DNA Technologies) and cloned into a modified pFL vector (MultiBac system, Geneva Biotech) resulting in an expression construct with an N-terminal melittin signal sequence followed by a FLAG-tag, His10-tag and a human rhinovirus 3C protease cleavage site. In the crystallisation construct, several modifications were introduced to the receptor to increase the functional expression yield, stability and crystallisation: To aid crystal contact formation in lipidic cubic phase 34 residues (R232-L265) of the ICL3 were replaced by the thermostable *Pyrococcus abysii* glycogen synthase (PGS) domain and 30 residues (360-389) were truncated from the C-terminus of the receptor. Analogous to previously reported structures, the gene was further modified by introducing mutations to improve protein yield and thermostability: A19T, R65H, V120L, A167V, Q193R, S322C, N325K and T333M were selected through two consecutive rounds of directed evolution in yeast^25^ on the wtOTR using the fluorescently labelled peptide antagonist PVA^63^ (HiLyte Fluor 647-Lys^8^ PVA (Eurogentec)), while D153A and S224A were identified to further increase thermostability as evidenced by an increase in melting temperature (Tm) in the CPM assay^49^.

For ligand-binding experiments, the codon-optimised wtOTR gene was subcloned into a mammalian expression vector (pcDNA3.1(+)) containing an N-terminal SNAP-tag (Cisbio). Point mutations were derived from this construct using SLIC cloning^64^.

### Expression and purification of OTR

Recombinant baculovirus was generated using the MultiBac expression system. The receptor expression was performed as previously described^30^. Briefly, *Spodoptera frugiperda* (*Sf*9) insect cells were infected with high-titer P1 or P2 baculovirus stocks (multiplicity of infection ≥5) at a cell density of 3 ×10^6^ cells/ml in Sf-900 II SFM medium (Thermo Fisher Scientific). Cells were harvested 72 h post-infection by centrifugation, washed with PBS, flash-frozen in liquid nitrogen and stored at −80 °C until further use.

Receptor-containing insect cell membranes were isolated by repeated Dounce homogenization in hypotonic (10 mM HEPES pH 7.5, 20 mM KCl, 10 mM MgCl_2_, 50 μg/ml Pefabloc SC (Carl Roth), 1 μg/ml Pepstatin A (Carl Roth)) and hypertonic buffer (10 mM HEPES pH 7.5, 20 mM KCl, 10 mM MgCl_2_, 1.0 M NaCl, 50 μg/ml Pefabloc SC, 1 μg/ml Pepstatin A) followed by a final wash in hypotonic buffer to reduce the NaCl concentration. The purified membrane fraction was resuspended in hypotonic buffer supplemented with 20 μM of retosiban, incubated for 30 min at 4 °C, frozen in liquid nitrogen and stored at −80°C until further use.

Prior to solubilisation, membranes were thawed on ice, retosiban was added to a final concentration of 100 μM and the suspension was incubated for 35 min in the presence of 2 mg/ml iodoacetamide (Sigma Aldrich) while turning. The membranes were then solubilised in 30 mM HEPES pH 7.5, 500 mM NaCl, 10 mM KCl, 5 mM MgCl_2_, 50 μg/ml Pefabloc SC, 1 μg/ml Pepstatin A, 1% (w/v) *n*-dodecyl-*β*-D-maltopyranoside (DDM, Anatrace) and 0.2% (w/v) cholesteryl hemisuccinate (CHS, Sigma Aldrich) at 4°C for 2.5 h. Insoluble material was removed by ultra-centrifugation and the cleared supernatant was incubated with TALON IMAC resin (GE Healthcare) overnight at 4°C.

The receptor-bound resin was washed with 30 column volumes (CV) of Wash Buffer I (50 mM HEPES pH 7.5, 500 mM NaCl, 10 mM MgCl_2_, 5 mM imidazole, 10% (v/v) glycerol, 1.0% (w/v) DDM, 0.2% (w/v) CHS, 8 mM ATP, 50 μM retosiban) followed by 30 CV of Wash Buffer II (50 mM HEPES pH 7.5, 500 mM NaCl, 15 mM imidazole, 10% (v/v) glycerol, 0.05% (w/v) DDM, 0.01% (w/v) CHS, 50 μM retosiban). Retosiban-bound OTR was eluted in fractions using a total of four CVs of Elution Buffer (50 mM HEPES pH 7.5, 500 mM NaCl, 250 mM imidazole, 10% (v/v) glycerol, 0.05% (w/v) DDM, 0.01% (w/v) CHS, 100 μM retosiban). Those fractions that contained purified OTR were concentrated to 0.5 ml using a 100 kDa MW cut-off Vivaspin 2 concentrator (Sartorius Stedim) and applied to a PD MiniTrap G-25 column (GE Healthcare) equilibrated with G25 Buffer (50 mM HEPES pH 7.5, 500 mM NaCl, 10% (v/v) glycerol, 0.03% (w/v) DDM, 0.006% (w/v) CHS, 100 μM retosiban) to remove imidazole. The eluted receptor-antagonist complex was treated overnight with Histagged 3C protease and PNGaseF (both prepared in-house) to remove the N-terminal affinity tags and deglycosylate the receptor. The mixture was then incubated with Ni-NTA resin (GE Healthcare) for 1 h, and cleaved receptor was collected as the flow-through. Purified OTR:retosiban complex was concentrated to ~70 mg/ml with a 100 kDa MW cut-off Vivaspin 2 concentrator. All protein concentrations were determined by measuring absorbance at 280 nm on a Nanodrop 2000 spectrophotometer (Thermo Fisher Scientific). For protein concentrations ?10 mg/ml 1:10 dilutions were prepared using the respective buffer to ensure measurement in the linear range of the instrument. Protein purity and monodispersity were assessed by SDS-PAGE and analytical size-exclusion chromatography using a Sepax Nanofilm SEC-250 column.

### Crystallisation in lipidic cubic phase

The OTR:retosiban complex was crystallised using the *in meso* method, however, at a reduced temperature of 16 °C. For reconstitution, the concentrated OTR:retosiban complex (~70 mg/ml) was mixed with molten monoolein (Sigma Aldrich) supplemented with 10% (w/w) cholesterol (Sigma Aldrich) using the twin-syringe method^65^. The final protein:lipid ratio was 40:60 (v/v). 40 nl boli were dispensed onto 96-well glass bases with a 120 μm spacer (Swissci), overlaid with 800 nl precipitant solution using a Gryphon LCP crystallisation robot (Art Robbins Instruments) and sealed with a cover glass. All initial steps were performed at 20 °C; however, for optimal crystal growth and diffraction the screening plates were subsequently transferred to a humidified incubator set to 16 °C. Optimised crystals used for data collection of retosiban-bound OTR were obtained in a condition consisting of 100 mM N-(2-acetamido)iminodiacetic acid (ADA) pH 5.8-6.0, 27-29% (v/v) PEG400, 320-350 mM NaH_2_PO_4_, 1 mM TCEP and 50 μM retosiban. Single crystals were mounted with 20 or 30 μm Dual-Thickness MicroMounts (MiTeGen) for data collection and cryo-cooled in liquid nitrogen without the addition of further cryoprotectant.

### Data collection, structure determination and refinement

X-ray diffraction data were collected using an EIGER 16M detector and a beam-size of 10×10 μm at the Swiss Light Source (SLS) of the Paul Scherrer Institute (PSI, Villigen, Switzerland). Datasets were collected using a beam attenuated to 30%, using 0.1° of oscillation and 0.05 s exposure time per frame. Data from individual crystals were integrated using *XDS*. Data from the nine best diffracting crystals were merged and scaled using the program *AIMLESS* from the CCP4 suite^66,67^. Data collection statistics are reported in **Table 1**.

Initial phase information was obtained by molecular replacement (MR) with the program *Phaser^6^* using a truncated poly-alanine model of the OX1R transmembrane domain (PDB ID 4ZJ8) and the separated PGS fusion protein^26^ as independent search models looking for one copy of each domain. Manual model building was performed in *COOT*^69^ using sigma-A weighted 2m|F_o_|-|DF_c_|, m|F_o_|-D|F_c_| maps together with simulated-annealing and simple composite omit maps calculated using *Phenix*^70^. Initial refinement was carried out with *REFMAC5^71^* using maximum-likelihood restrained refinement in combination with the jellybody protocol. Further and final stages of refinement were performed with *Phenix.refine*^72^ with positional, individual isotropic B-factor refinement and *TLS*. The final refinement statistics are presented in **Table 1**. Co-ordinates and structure factors have been deposited in the Protein Data Bank under accession code 6TPK.

### Whole-cell ligand binding assay

HEK293T/17 cells (ATCC) were cultivated in Dulbecco’s modified medium (Sigma) supplemented with 100 units/ml penicillin, 100 μg/ml streptomycin (Sigma) and 10% (v/v) fetal calf serum (BioConcept). Cells were maintained at 37°C in a humidified atmosphere of 5% CO_2_, 95% air. Transient transfections were performed with TransIT-293 (Mirus Bio) according to the manufacturer’s instructions.

Ligand binding experiments were performed on whole HEK293T cells for comparison of affinities for wild-type and receptor mutants using a homogeneous time-resolved fluorescence (HTRF) binding assay. HEK293T cells were seeded and transfected in poly-D-lysine-coated 384-well plates (Greiner) at a cell density of 7500 cells per well. 48 h after transfection, cells were labelled with 50 nM SNAP-Lumi4-Tb (Cisbio) in assay buffer (20 mM HEPES pH 7.5, 100 mM NaCl, 3 mM MgCl_2_ and 0.05% (w/v) BSA) for 1.5 h at 37°C. Cells were washed four times with assay buffer and were then incubated for 1 h at RT in assay buffer containing fluorescently labelled tracer peptide (HiLyte Fluor 647-Lys^8^ PVA). Depending on the determined KD from saturation binding at each mutant, tracer peptide for competition binding was used at concentrations of 5, 25 or 150 nM together with a concentration range of unlabelled retosiban as competitor. For characterisation of the Mg^2+^-binding site and its influence on oxytocin-binding, fluorescently labelled oxytocin (HiLyte Fluor 488-Orn^8^ oxytocin (Eurogentec)) was used. Cells were washed in wash buffer (20 mM HEPES pH 7.5, 100 mM KCl and 0.05% (w/v) BSA) and assay buffer (20 mM HEPES pH 7.5, 100 mM KCl, 3 mM MgCl_2_ or 2 mM EDTA and 0.05% (w/v) BSA) was adjusted to contain either divalent magnesium or the chelator EDTA. Fluorescence intensities were measured on an Infinite M1000 fluorescence plate reader (Tecan) with an excitation wavelength of 340 nm. Acceptor emission was measured at 520 nm or 665 nm for the HiLyte Fluor 488 or 647, respectively. The Tb^3+^ donor emission was recorded at 620 nm. The ratio of FRET-donor and acceptor fluorescence intensities was calculated (F665 nm/F520 nm or F620 nm). Total binding was obtained in the absence of competitor, and nonspecific binding was determined in the presence of 1000-fold excess of unlabelled competitor. Data were normalised to the specific binding for each individual experiment and were analysed by global fitting to a one-site heterologous competition equation with the GraphPad Prism software (version 8.1.1, GraphPad Prism). To obtain Ki values, data were corrected for fluorescent ligand occupancy of each mutant with the Cheng-Prusoff equation as K_i_ = IC_50_/(1 + [fl. ligand] / K_*d*_).

### Data availability

Atomic coordinates and structure factors of the OTR:retosiban complex structure have been deposited in the Protein Data Bank under accession code 6TPK. Data supporting the findings of this manuscript are available from the corresponding author upon reasonable request.

## Acknowledgements

We thank B. Blattmann of the Protein Crystallisation Center at the University of Zurich for his support with crystallisation, the staff of the X06SA beamline at the Paul Scherrer Institute for support during data collection and I. Berger at the European Molecular Biology Laboratory for providing us with baculovirus transfer vectors. We further thank C. Thom for help with protein expression, L. Wiedmer for helpful discussions during structure analysis and GlaxoSmithKline for the generous gift of retosiban. This work was supported by Schweizerischer Nationalfonds Grants 31003A_153143 and 31003A_182334, both to A.P..

## Author contributions

Y.W. and L.K. created libraries and performed directed evolution in yeast. Y.W. and J.S. carried out the mutagenesis and thermostabilisation of the receptor and designed and characterised crystallisation constructs. Y.W. and J.E. expressed the protein. Y.W. and J.S. purified and crystallised the OTR-PGS fusion protein and harvested crystals. J.E. supported cloning and expression. Y.W., J.S. and J.E collected and processed the data, and solved and refined the structures. Y.W. performed ligand-binding experiments and analysed the data. Project management was carried out by Y.W. and A.P. The manuscript was prepared by Y.W., J.S., J.E., and A.P. All authors contributed to the final editing and approval of the manuscript.

## Competing interests

The authors declare no competing interests.

## References

1. Donaldson, Z.R. & Young, L.J. Oxytocin, vasopressin, and the neurogenetics of sociality. Science 322, 900–904 (2008).

2. Kimura, T., Tanizawa, O., Mori, K., Brownstein, M.J. & Okayama, H. Structure and expression of a human oxytocin receptor. Nature 356, 526–529 (1992).

3. Barberis, C., Mouillac, B. & Durroux, T. Structural bases of vasopressin/oxytocin receptor function. J. Endocrinol. 156, 223–229 (1998).

4. Duvigneaud, V., Lawler, H.C. & Popenoe, E.A. Enzymatic cleavage of glycinamide from vasopressin and a proposed structure for this pressor-antidiuretic hormone of the posterior pituitary. J. Am. Chem. Soc. 75, 4880–4881 (1953).

5. Duvigneaud, V., Ressler, C. & Trippett, S. The sequence of amino acids in oxytocin, with a proposal for the structure of oxytocin. J. Biol. Chem. 205, 949–957 (1953).

6. Carter, C.S. Neuroendocrine perspectives on social attachment and love. Psychoneuroendocrinology 23, 779–818 (1998).

7. Insel, T.R. A neurobiological basis of social attachment. Am. J. Psychiatry 154, 726735 (1997).

8. Dale, H.H. On some physiological actions of ergot. J. Physiol. 34, 163–206 (1906).

9. Fuchs, A.R., Fuchs, F., Husslein, P., Soloff, M.S. & Fernstrom, M.J. Oxytocin receptors and human parturition - a dual role for oxytocin in the Initiation of labor. Science 215, 1396–1398 (1982).

10. Uvnäs-Moberg, K., Widstrom, A.M., Werner, S., Matthiesen, A.S. & Winberg, J. Oxytocin and prolactin levels in breast-feeding women. Correlation with milk yield and duration of breast-feeding. Acta Obstet. Gynecol. Scand. 69, 301–306 (1990).

11. Ott, I. & Scott, J.C. The galactagogue action of the thymus and corpus luteum. Exp. Biol. Med. 8, 49–49 (1910).

12. Modahl, C., Fein, D., Waterhouse, L. & Newton, N. Does oxytocin deficiency mediate social deficits in autism? J. Autism Dev. Disord. 22, 449–451 (1992).

13. Andari, E. et al. Promoting social behavior with oxytocin in high-functioning autism spectrum disorders. Proc. Natl. Acad. Sci. U. S. A. 107, 4389–4394 (2010).

14. Hollander, E. et al. Oxytocin infusion reduces repetitive behaviors in adults with autistic and Asperger’s disorders. Neuropsychopharmacology 28, 193–198 (2003).

15. McCarthy, M.M. & Altemus, M. Central nervous system actions of oxytocin and modulation of behavior in humans. Mol. Med. Today 3, 269–275 (1997).

16. Pedersen, C.A. et al. Intranasal oxytocin reduces psychotic symptoms and improves Theory of Mind and social perception in schizophrenia. Schizophr. Res. 132, 50–53 (2011).

17. Windle, R.J., Shanks, N., Lightman, S.L. & Ingram, C.D. Central oxytocin administration reduces stress-induced corticosterone release and anxiety behavior in rats. Endocrinology 138, 2829–2834 (1997).

18. Giuliano, F. & Clèment, P. Pharmacology for the treatment of premature ejaculation. Pharmacol. Rev 64, 621–644 (2012).

19. Moraloglu, O., Tonguc, E., Var, T., Zeyrek, T. & Batioglu, S. Treatment with oxytocin antagonists before embryo transfer may increase implantation rates after IVF. Reprod. Biomed. Online 21, 338–343 (2010).

20. Andersen, L.F., Lyndrup, J., Åkerlund, M. & Melin, P. Oxytocin receptor blockade: a new principle in the treatment of preterm labor? Am. J. Perinatol. 6, 196–199 (1989).

21. Theobald, G.W., Graham, A., Campbell, J., Gange, P.D. & Driscoll, W.J. The use of post-pituitary extract in physiological amounts in obstetrics - a preliminary report. BMJ 2, 123–127 (1948).

22. Tsatsaris, V., Carbonne, B. & Cabrol, D. Atosiban for preterm labour. Drugs 64, 375382 (2004).

23. Klein, U., Gimpl, G. & Fahrenholz, F. Alteration of the myometrial plasma membrane cholesterol content with beta-cyclodextrin modulates the binding affinity of the oxytocin receptor. Biochemistry 34, 13784–13793 (1995).

24. Soloff, M.S. & Swartz, T.L. Characterization of a proposed oxytocin receptor in rat mammary-gland. J. Mol. Biol. 248, 6471–6478 (1973).

25. Schütz, M. et al. Directed evolution of G protein-coupled receptors in yeast for higher functional production in eukaryotic expression hosts. Sci. Rep. 6, 21508 (2016).

26. Yin, J., Mobarec, J.C., Kolb, P. & Rosenbaum, D.M. Crystal structure of the human OX2 orexin receptor bound to the insomnia drug suvorexant. Nature 519, 247–250 (2015).

27. Ballesteros, J.A. & Weinstein, H. [19] Integrated methods for the construction of threedimensional models and computational probing of structure-function relations in G protein-coupled receptors. in Methods in Neurosciences, Vol. 25 (ed. Sealfon, S.C.) 366–428 (Academic Press, 1995).

28. Manglik, A. et al. Crystal structure of the micro-opioid receptor bound to a morphinan antagonist. Nature 485, 321–326 (2012).

29. Miller-Gallacher, J.L. et al. The 2.1 A resolution structure of cyanopindolol-bound beta1-adrenoceptor identifies an intramembrane Na+ ion that stabilises the ligand-free receptor. PLoS One 9, e92727 (2014).

30. Schöppe, J. et al. Crystal structures of the human neurokinin 1 receptor in complex with clinically used antagonists. Nat. Commun. 10, 17 (2019).

31. White, J.F. et al. Structure of the agonist-bound neurotensin receptor. Nature 490, 508513 (2012).

32. Yang, Z. et al. Structural basis of ligand binding modes at the neuropeptide Y Y1 receptor. Nature 556, 520–524 (2018).

33. Warne, T., Edwards, P.C., Dore, A.S., Leslie, A.G.W. & Tate, C.G. Molecular basis for high-affinity agonist binding in GPCRs. Science 364, 775–778 (2019).

34. Draper-Joyce, C.J. et al. Structure of the adenosine-bound human adenosine A1 receptor-Gi complex. Nature 558, 559–563 (2018).

35. Borthwick, A.D. & Liddle, J. The design of orally bioavailable 2, 5-diketopiperazine oxytocin antagonists: from concept to clinical candidate for premature labor. Med. Res. Rev. 31, 576–604 (2011).

36. Liddle, J. et al. The discovery of GSK221149A: a potent and selective oxytocin antagonist. Bioorg. Med. Chem. Lett. 18, 90–94 (2008).

37. Wyatt, P.G. et al. 2,5-Diketopiperazines as potent and selective oxytocin antagonists 1: Identification, stereochemistry and initial SAR. Bioorg. Med. Chem. Lett. 15, 2579–2582 (2005).

38. Mouillac, B. et al. The binding site of neuropeptide vasopressin V1a receptor: evidence for a major localization within transmembrane regions. J. Biol. Chem. 270, 25771–25777 (1995).

39. Hawtin, S.R., Ha, S.N., Pettibone, D.J. & Wheatley, M. A Gly/Ala switch contributes to high affinity binding of benzoxazinone-based non-peptide oxytocin receptor antagonists. FEBS Lett. 579, 349–356 (2005).

40. Burger, K., Gimpl, G. & Fahrenholz, F. Regulation of receptor function by cholesterol. Cell. Mol. Life. Sci. 57, 1577–1592 (2000).

41. Hulce, J.J., Cognetta, A.B., Niphakis, M.J., Tully, S.E. & Cravatt, B.F. Proteome-wide mapping of cholesterol-interacting proteins in mammalian cells. Nat. Methods 10, 259264 (2013).

42. Lingwood, D. & Simons, K. Lipid rafts as a membrane-organizing principle. Science 327, 46–50 (2010).

43. Gimpl, G., Burger, K. & Fahrenholz, F. Cholesterol as modulator of receptor function. Biochemistry 36, 10959–10974 (1997).

44. Paila, Y.D. & Chattopadhyay, A. Membrane cholesterol in the function and organization of G-protein coupled receptors. Subcell. Biochem. 51, 439–466 (2010).

45. Cherezov, V. et al. High-resolution crystal structure of an engineered human beta2-adrenergic G protein-coupled receptor. Science 318, 1258–1265 (2007).

46. Zhang, K. et al. Structure of the human P2Y12 receptor in complex with an antithrombotic drug. Nature 509, 115–118 (2014).

47. Gimpl, G. Interaction of G protein coupled receptors and cholesterol. Chem. Phys. Lipids 199, 61–73 (2016).

48. Gimpl, G. & Fahrenholz, F. Cholesterol as stabilizer of the oxytocin receptor. BBA-Biomembranes 1564, 384–392 (2002).

49. Alexandrov, A.I., Mileni, M., Chien, E.Y., Hanson, M.A. & Stevens, R.C. Microscale fluorescent thermal stability assay for membrane proteins. Structure 16, 351–359 (2008).

50. Yao, Z. & Kobilka, B. Using synthetic lipids to stabilize purified β2 adrenoceptor in detergent micelles. Anal. Biochem. 343, 344–346 (2005).

51. Hanson, M.A. et al. A specific cholesterol binding site is established by the 2.8 angstrom structure of the human beta(2)-adrenergic receptor. Structure 16, 897–905 (2008).

52. Thompson, A.A. et al. GPCR stabilization using the bicelle-like architecture of mixed sterol-detergent micelles. Methods 55, 310–317 (2011).

53. Vukoti, K., Kimura, T., Macke, L., Gawrisch, K. & Yeliseev, A. Stabilization of functional recombinant cannabinoid receptor CB2 in detergent micelles and lipid bilayers. PLOS ONE 7, e46290 (2012).

54. Degorce, F. et al. HTRF: A technology tailored for drug discovery - a review of theoretical aspects and recent applications. Curr. Chem. Genomics 3, 22–32 (2009).

55. Postina, R., Kojro, E. & Fahrenholz, F. Separate agonist and peptide antagonist binding sites of the oxytocin receptor defined by their transfer into the V2 vasopressin receptor. J. Biol. Chem. 271, 31593–31601 (1996).

56. Katritch, V. et al. Allosteric sodium in class A GPCR signaling. Trends Biochem. Sci. 39, 233–244 (2014).

57. Schiffmann, A. & Gimpl, G. Sodium functions as a negative allosteric modulator of the oxytocin receptor. BBA-Biomembranes 1860, 1301–1308 (2018).

58. Antoni, F.A. & Chadio, S.E. Essential role of magnesium in oxytocin-receptor affinity and ligand specificity. Biochem. J. 257, 611–614 (1989).

59. Pearlmutter, A.F. & Soloff, M.S. Characterization of the metal ion requirement for oxytocin-receptor interaction in rat mammary gland membranes. J. Biol. Chem. 254, 3899–3906 (1979).

60. Larivière, R. & Schiffrin, E.L. Effects of monovalent and divalent cations and of guanine nucleotides on binding of vasopressin to the rat mesenteric vasculature. Canadian Journal of Physiology and Pharmacology 65, 1171–1181 (1987).

61. Wootten, D.L., Simms, J., Massoura, A.J., Trim, J.E. & Wheatley, M. Agonist-specific requirement for a glutamate in transmembrane helix 1 of the oxytocin receptor. Mol. Cell. Endocrinol 333, 20–27 (2011).

62. Rodrigo, J. et al. Mapping the binding site of arginine vasopressin to V1a and V1b vasopressin receptors. Mol. Endocrinol. 21, 512–523 (2007).

63. Mouillac, B., Manning, M. & Durroux, T. Fluorescent agonists and antagonists for vasopressin/oxytocin G protein-coupled receptors: usefulness in ligand screening assays and receptor studies. Mini Rev. Med. Chem. 8, 996–1005 (2008).

64. Li, M.Z. & Elledge, S.J. SLIC: A Method for Sequence- and Ligation-Independent Cloning. in Gene Synthesis: Methods and Protocols (ed. Peccoud, J.) 51–59 (Humana Press, Totowa, NJ, 2012).

65. Caffrey, M. & Cherezov, V. Crystallizing membrane proteins using lipidic mesophases. Nat. Protoc. 4, 706–731 (2009).

66. Dodson, E.J., Winn, M. & Ralph, A. Collaborative computational project number 4 “the CCP4 suite: programs for protein crystallography”. Acta Crystallogr. D 50, 760–763 (1994).

67. Evans, P.R. & Murshudov, G.N. How good are my data and what is the resolution? Acta Crystallogr. D 69, 1204–1214 (2013).

68. McCoy, A.J. et al. Phaser crystallographic software. J. Appl. Crystallogr. 40, 658–674 (2007).

69. Emsley, P., Lohkamp, B., Scott, W.G. & Cowtan, K. Features and development of Coot. Acta Crystallogr. D 66, 486–501 (2010).

70. Adams, P.D. et al. PHENIX: a comprehensive Python-based system for macromolecular structure solution. Acta Crystallogr. D 66, 213–221 (2010).

71. Murshudov, G.N. et al. REFMAC5 for the refinement of macromolecular crystal structures. Acta Crystallogr. D 67, 355–367 (2011).

72. Afonine, P.V. et al. Towards automated crystallographic structure refinement with phenix.refine. Acta Crystallogr. D 68, 352–367 (2012).

