## Supplementary for "Crystal structure of the human oxytocin receptor"

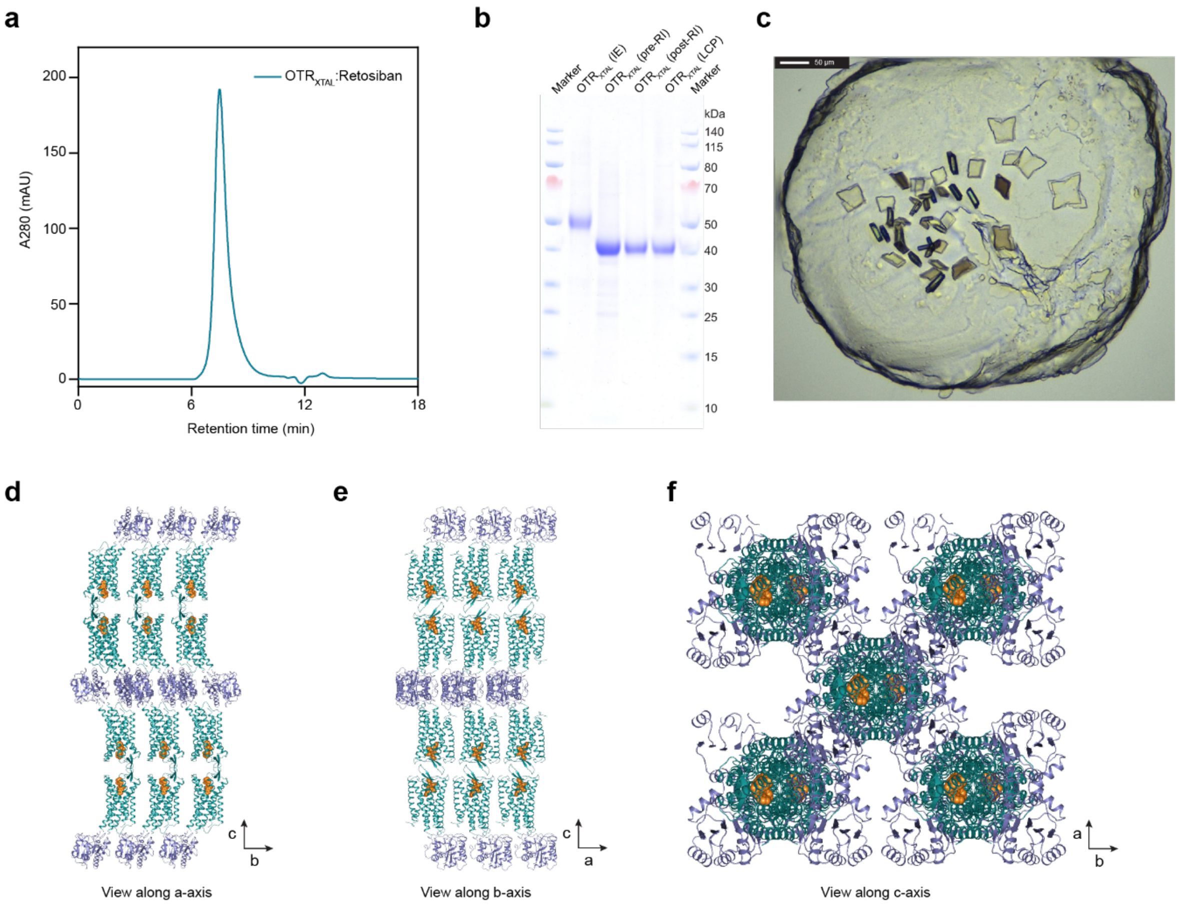


**Supplementary Fig. 1 | OTR crystallisation**

**a**, Size exclusion profile of purified OTR:retosiban complex. **b**, SDS-PAGE gel of purified OTR:retosiban complex. Loading scheme from left to right: IMAC eluate (IE), IMAC eluate after treatment with 3C protease and PNGaseF (pre-RI), reverse IMAC eluate (RI), concentrated reverse IMAC eluate used for crystallisation (LCP). **c**, Bright-field image of OTR:retosiban crystals in lipidic cubic phase. **d**, Packing of OTR:retosiban crystals as viewed along the a axis of the unit cell. **e**, Packing of OTR:retosiban crystals as viewed along the b axis of the unit cell. **f**, Packing of OTR:retosiban crystals as viewed along the c axis of the unit cell.

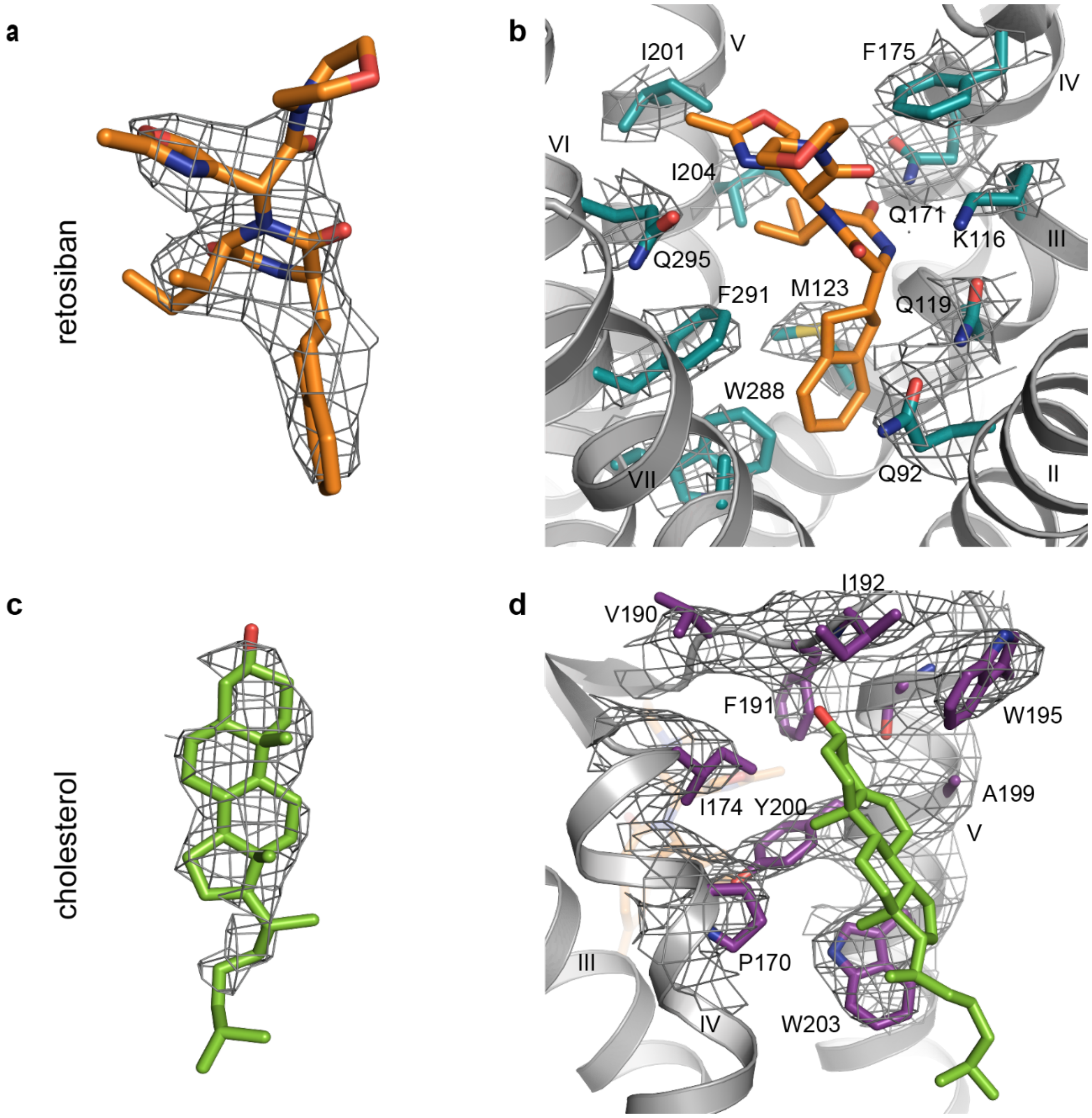


**Supplementary Fig. 2 | Electron density maps of the extracellular ligand binding pocket and the cholesterol binding site of OTR**

**a**, Retosiban with 2Fo-Fc electron density map contoured at 1.0 σ. **b**, 2Fo-Fc electron density map contoured at 1.0 σ for OTR residues interacting with retosiban. Oxygen and nitrogen atoms are coloured in red and blue, respectively. **c**, Cholesterol with 2Fo-Fc electron density map contoured at 1.0 σ. **d**, 2Fo-Fc electron density map contoured at 1.0 σ for OTR residues interacting with cholesterol.

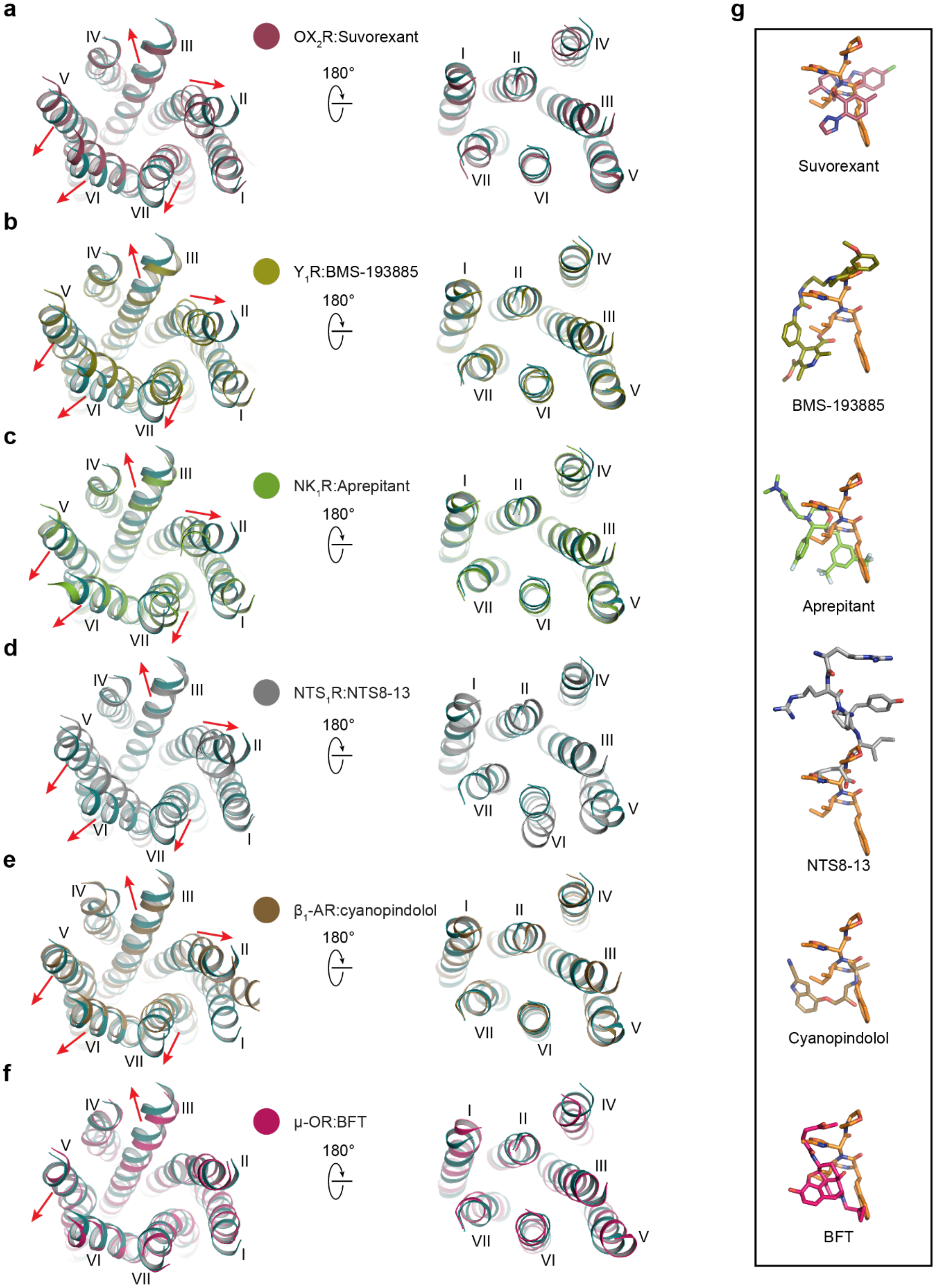
a
IV
III
II
V
VI
VII
I
OX2R:Suvorexant
180°
b
Y1R:BMS-193885
c
NK1R:Aprepitant
d
NTS1R:NTS8-13
e
β1-AR:cyanopindolol
f
μ-OR:BFT
g
Suvorexant
BMS-193885
Aprepitant
NTS8-13
Cyanopindolol
BFT

**Supplementary Fig. 3 | Comparison of transmembrane helix conformations shaping the extracellular binding site of OTR to other peptidergic and aminergic GPCRs**

As viewed from the extracellular space (left) and from the cytoplasm (right). Red arrows indicate shift of OTR extracellular helix tips relative to reference receptor.

**a-f**, Structural superposition of OTR:retosiban (coloured in cyan) with previously reported class A GPCR crystal structures yields root-mean-square deviations (RMSD, summarised in Supplementary Table 1) for backbone atoms of (**a**) 2.7 Å to orexin 2 receptor, (**b**) 2.9 Å to neuropeptide Y Y1 receptor, (**c**) 1.6 Å to neurokinin 1 receptor, (**d**) 2.3 Å to neurotensin 1 receptor, (**e**) 2.6 Å to $\beta_1$-adrenergic receptor and (**f**) 2.6 Å to the $\mu$-opioid receptor.

(**g**) Isolated ligands from the overlays in (**a-f**), as viewed from helix VI-VII (retosiban coloured in orange).

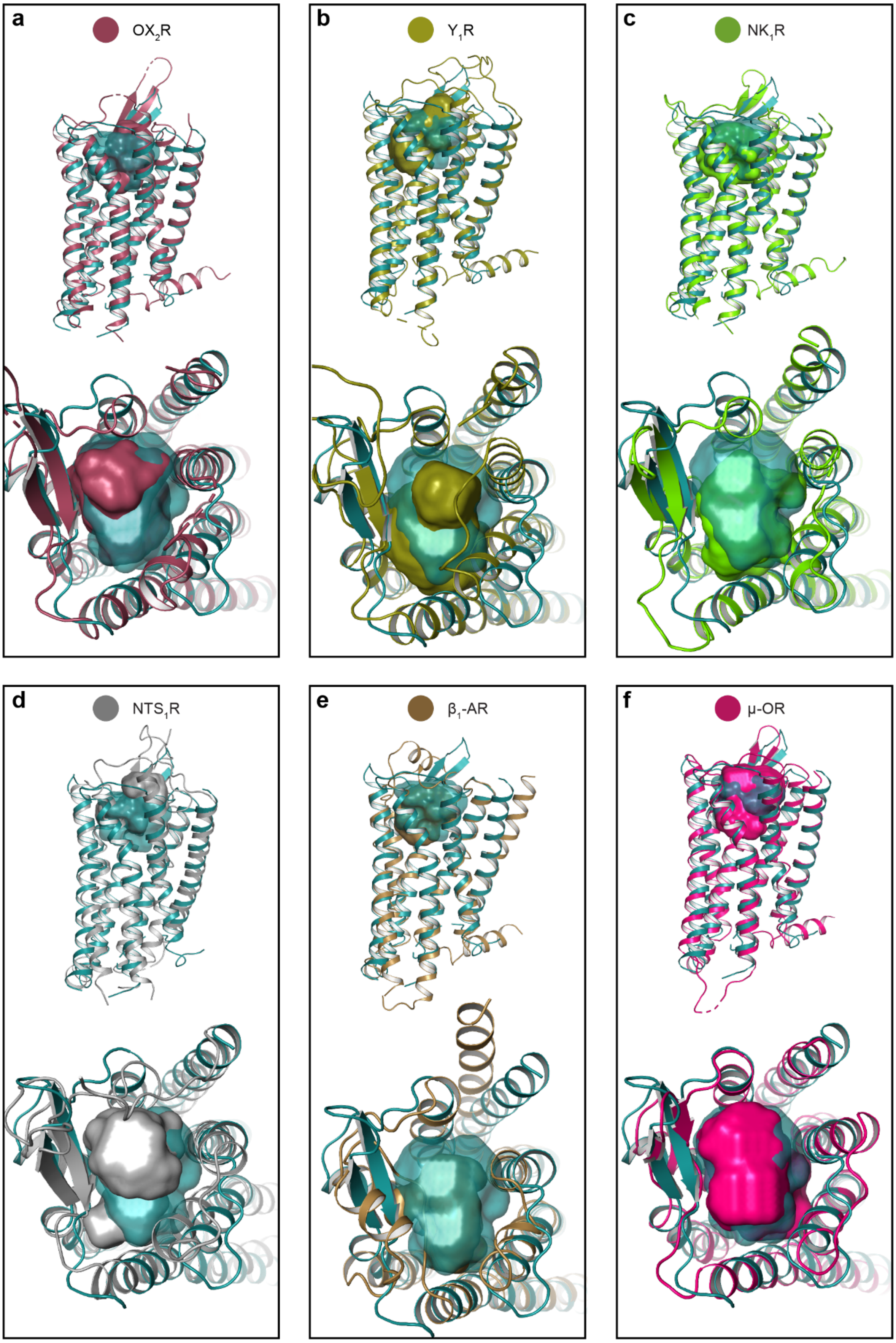

a
OX2R
b
Y1R
c
NK1R
d
NTS1R
e
β1-AR
f
μ-OR

**Supplementary Fig. 4 | Volume differences of the orthosteric binding pocket between OTR and other peptidergic and aminergic GPCRs**

**a-f**, In each panel an overlay of OTR:retosiban (coloured in cyan) with another GPCR is shown in cartoon representation, as viewed from the membrane plane (top) and from the extracellular space (bottom). The volumes of the extracellular ligand binding pockets are outlined as solid surfaces and highlighted in the respective colour of the receptors. Volumes were calculated using the program POVME 2.0 (Durrant, J.D. *et al*. *J. Chem. Theory Comput.* **10**, 5047-5056 (2014)). The structures overlayed with OTR:retosiban are (**a**) $OX_2R$:Suvorexant (PDB ID: 4S0V), (**b**) $Y_1R$:BMS-193885 (PDB ID: 5ZBH), (**c**) $NK_1R$:Aprepitant (PDB ID: 6HLO), (**d**) $NTS_1R$:NTS8-13 (PDB ID: 4GRV), (**e**) $\beta_1$ AR:cyanopindolol (PDB ID: 4BVN) and (**f**) μ-OR:BFT (PDB ID: 4DKL).

**a**

| Group | Species | Accession | 1.34 | 1.35 | 1.36 | 2.64 | 2.65 | 2.66 | 5.37 | 5.38 | 5.39 | 5.40 | 5.41 | 5.42 | |
|---|---|---|---|---|---|---|---|---|---|---|---|---|---|---|---|
| Mam | Human (homo sapiens sapiens) | P30559 | V | E | V | W | D | I | A | Y | I | T | W | I | OTR |
| | Gorilla (Gorilla gorilla gorilla) | G3S6I2 | V | E | V | W | D | I | A | Y | I | T | W | I | |
| | Bovine (Bos taurus) | P56449 | V | E | V | W | D | I | A | Y | I | T | W | I | |
| | Mouse (mus musculus) | P97926 | V | E | V | W | D | I | A | Y | V | T | W | I | |
| | Human (homo sapiens sapiens) | P37288 | L | E | I | W | D | I | A | Y | V | T | W | M | $V_{1A}R$ |
| | Gorilla (Gorilla gorilla gorilla) | XP_018893576.1# | L | E | I | W | D | I | A | Y | V | T | W | M | |
| | Bovine (Bos taurus) | A2VDS9 | L | E | I | W | D | I | A | Y | V | T | W | M | |
| | Mouse (mus musculus) | Q62463 | L | E | V | W | D | I | A | Y | V | T | W | M | |
| Non-mammal vertebrates | Turkey (Meleagris gallogavo) | G3USG7 | V | E | V | W | D | I | A | Y | V | T | W | I | Homologues |
| | Zebrafish (Danio rerio) | E4W699 | V | E | V | W | D | I | A | Y | I | T | W | I | |
| | King cobra (Ophiophagus hannah) | V8P3C6 | V | E | V | W | D | I | A | Y | V | T | W | M | |
| | Coelacanth (Latimeria chalumnae) | XP_005988002.1# | V | E | V | W | D | I | A | Y | V | T | W | I | |
| | Xenopus (Xenopus laevis) | XP_018116839.1# | V | E | V | W | D | I | T | Y | I | T | W | I | |
| Invertebrates | Snail (Lymnaea stagnalis) | AAA91998.1# | I | E | I | S | D | V | A | Y | I | T | W | V | |
| | Ant (Harpegnathos saltator) | XP_011151734.1# | W | E | I | W | D | I | A | Y | I | T | W | Y | |
| | Beetle (Tribolium castaneum) | NP_001078830.1# | V | E | V | W | D | I | A | Y | V | T | W | Y | |

**b**

| Receptor | Accession | 1.34 | 1.35 | 1.36 | 2.64 | 2.65 | 2.66 | 5.37 | 5.38 | 5.39 | 5.40 | 5.41 | 5.42 |
|---|---|---|---|---|---|---|---|---|---|---|---|---|---|
| OTR | P30559 | V | E | V | W | D | I | A | Y | I | T | W | I |
| $OX_2R$ | O43614 | V | L | I | D | I | T | M | Y | H | I | C | F |
| $Y_1R$ | P25929 | T | L | A | T | L | M | S | Y | T | T | L | L |
| $NK_1R$ | P25103 | L | W | A | Y | A | V | V | Y | H | I | C | V |
| $NTS_1R$ | P30989 | L | V | T | F | I | W | V | V | I | Q | V | N |
| $\beta_1$ | P08588 | G | M | G | V | V | W | A | Y | A | I | A | S |
| μ-OR | P35372 | T | I | M | Y | L | M | L | L | K | I | C | V |

**Supplementary Fig. 5 | Alignment of the divalent cation binding site of OTR**

**a**, Amino acid alignment of residues compromising the human OTR divalent cation coordination (red) and cholesterol binding (green) sites and several vertebrate and invertebrate homologues. Species name and UniProt Knowledgebase accession codes (www.uniprot.org) or NCBI reference sequence (marked with #, www.ncbi.nlm.nih.gov) are indicated in the first and second column, respectively. Amino acid residue alignment of positions according to the Ballesteros-Weinstein numbering scheme (BW) are indicated above the sequences. Residues of the OTR magnesium binding site are highlighted by black boxes. Acidic residues are highlighted in red. **b**, Alignment with other peptidergic and aminergic GPCRs. Receptor name and UniProt Knowledgebase (www.uniprot.org) accession codes are indicated in the first and second column, respectively. Amino acid residue alignment of positions 1.34-1.36 and 2.64-2.66 according to the Ballesteros-Weinstein numbering scheme (BW) are indicated above the sequences. Residues of the OTR magnesium binding site are highlighted by black boxes. Acidic residues are highlighted in red.

**Supplementary Table 1 | Extracellular ligand binding pocket comparison.**

| receptor | RMSD [Å] | ligand | $M_w$ [g mol$^{-1}$] | pocket volume [Å$^3$] | pocket volume relative to OTR [%] | PDB ID |
|---|---|---|---|---|---|---|
| OTR | | retosiban | 494.6 | 792 | 100 | xx |
| OX2R | 2.7 | Suvorexant | 450.9 | 569 | 72 | 4S0V |
| $Y_1R$ | 2.9 | BMS-193885 | 590.7 | 605 | 76 | 5ZBH |
| $NK_1R$ | 1.6 | Aprepitant | 534.4 | 597 | 75 | 6HLO |
| $NTS_1R$ | 2.3 | NTS8-13 | 817.0 | 433 | 55 | 4GRV |
| $\beta_1$-AR | 2.6 | Cyanopindolol | 287.4 | 320 | 40 | 4BVN |
| μ-OR | 2.6 | BFT | 468.5 | 911 | 115 | 4DKL |

Root-mean-square deviations for backbone atoms (RMSD). Pocket volumes were calculated using the program POVME 2.0 (Durrant, J.D. *et al*. *J. Chem. Theory Comput.* **10**, 5047-5056 (2014)).

**Supplementary Table 2 | Binding of retosiban to wild-type and mutated OTR variants.**

| construct | $K_i$ [nM] | 95% CI [nM] |
|---|---|---|
| Q92A | 16.3 | 11.7 - 22.7 |
| Q92N | 517.6 | 411.1 - 650.8 |
| K116A | 195.4 | 158.4 - 241.3 |
| Q119A | 2.2 | 1.7 - 3.0 |
| Q119N | 31.1 | 11.1 - 85.4 |
| Q119E | 1142.9 | 767.3 - 1713 |
| M123A | n.b. | |
| Q171A | 8260.4 | 5221 - 13060 |
| F175A | 11.7 | 8.9 - 15.3 |
| I201A | n.b. | |
| I204A | n.b. | |
| F291A | 1078.9 | 897.1 - 1299 |
| A318G | 40.5 | 25.2 - 65.3 |
| wtOTR | 2.6 | 2.0 - 3.3 |
| $OTR_{XTAL}$ | 7.6 | 5.9 - 9.9 |

Whole-cell competition binding experiments of different concentrations of unlabelled retosiban to HEK293T cells expressing wild-type (wtOTR) and mutated wtOTR variants, in the presence of fluorescently labelled PVA as a competitor. Data are shown as mean values ($K_i$) with the respective 95% confidence intervals (CI) from 3-5 independent experiments performed in duplicate. *n.b.*, no binding. $K_i$ values were calculated using the Cheng-Prusoff equation (Cheng, Y. & Prusoff, W.H. *Biochem. Pharmacol.* **22**, 3099-108 (1973)).

**Supplementary Table 3 | Binding of agonist or antagonist to wild-type OTR and variants with mutations in the cholesterol binding site.**

| construct | surface expression (% of wtOTR) | $B_{max}$ agonist (% of wtOTR) | $B_{max}$ antagonist (% of wtOTR) |
|---|---|---|---|
| Y200A | 111.3 ± 17.1 | 4.6 ± 4.1 | 0.7 ± 0.03 |
| Y200H | 108.0 ± 13.3 | 11.7 ± 1.5 | 3 ± 0.4 |
| W203A | 104.5 ± 16.5 | 19.2 ± 3.5 | 8.1 ± 0.5 |
| W203H | 101.3 ± 27.5 | 5.1 ± 1.5 | 0.2 ± 0.02 |
| wt | 100 ± 13.9 | 100 ± 20.5 | 100 ± 3.5 |
| $OTR_{XTAL}$ | 182.7 ± 38.5 | 471.4 ± 50.4 | 419.7 ± 44.2 |

Whole-cell specific binding experiments of agonist (HL488-Orn$^{8}$-OT) and antagonist (HL647-PVA) on HEK293T cells expressing wild-type (wtOTR) and mutated wtOTR variants expressed as $B_{max}$ values. In comparison the surface expression is also shown. Data are shown as mean values (surface expression and $B_{max}$) ± standard deviations (SD) from 3 independent experiments performed in triplicates.

**Supplementary Table 4 | Thermostability of $OTR_{XTAL}$ and variants with mutations in the cholesterol binding site.**

| construct | $T_m$ + CHS [°C] | $T_m$ - CHS [°C] |
|---|---|---|
| $OTR_{XTAL}$ Y200A | 55.2 ± 0.8 | 44.5 ± 3.9 |
| $OTR_{XTAL}$ W203A | 61.2 ± 0.2 | 41.9 ± 3.1 |
| $OTR_{XTAL}$ | 64.2 ± 0.6 | 43.7 ±1.9 |

CPM thermostability assay of $OTR_{XTAL}$ and mutated variants thereof in the presence or absence of the cholesterol analogue cholesteryl hemisuccinate (CHS). Data are shown as mean values ($T_m$) ± standard deviations (SD) from 3 independent experiments.

**Supplementary Table 5 | Ligand binding to wild-type OTR and variants with mutations in the $Mg^{2+}$ binding site.**

| construct | agonist | antagnonist | $Mg^{2+}$ | $K_D$ [nM] | 95% CI [nM] |
|---|---|---|---|---|---|
| wtOTR | + | | + | 1.2 | 0.9 - 1.5 |
| wtOTR | + | | - | 27.9 | 22.7 - 34.4 |
| E42A | + | | + | 19.1 | 15 - 24.4 |
| E42A | + | | - | 29.6 | 23 - 38.4 |
| D100A | + | | + | 26.2 | 23.8 - 28.8 |
| D100A | + | | - | 30.8 | 28.1 - 33.7 |
| E42D, D100E | + | | + | 16.4 | 11.2 - 24.1 |
| E42D, D100E | + | | - | 28.4 | 20.1 - 40.7 |

Whole-cell specific binding experiment of agonist (HL488-Orn$^8$-OT) or antagonist (HL647-PVA) to HEK293T cells expressing wild-type (wtOTR) and mutated wtOTR variants in the presence (+) or absence (-) of 3 mM $Mg^{2+}$. Data are shown as mean values ($K_D$) with the respective 95 % confidence interval (CI) from three independent experiments performed in triplicates.
